# The projection of a test genome onto a reference population and applications to humans and archaic hominins

**DOI:** 10.1101/008805

**Authors:** Melinda A. Yang, Kelley Harris, Montgomery Slatkin

## Abstract

We introduce a method for comparing a test genome with numerous genomes from a reference population. Sites in the test genome are given a weight *w* that depends on the allele frequency *x* in the reference population. The projection of the test genome onto the reference population is the average weight for each *x*, 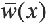. The weight is assigned in such a way that if the test genome is a random sample from the reference population, 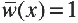. Using analytic theory, numerical analysis, and simulations, we show how the projection depends on the time of population splitting, the history of admixture and changes in past population size. The projection is sensitive to small amounts of past admixture, the direction of admixture and admixture from a population not sampled (a ghost population). We compute the projection of several human and two archaic genomes onto three reference populations from the 1000 Genomes project, Europeans (CEU), Han Chinese (CHB) and Yoruba (YRI) and discuss the consistency of our analysis with previously published results for European and Yoruba demographic history. Including higher amounts of admixture between Europeans and Yoruba soon after their separation and low amounts of admixture more recently can resolve discrepancies between the projections and demographic inferences from some previous studies.

## Introduction

The wealth of genomic data now available calls for new methods of analysis. One class of methods estimates parameters of demographic models using samples from multiple populations. Such methods are computationally challenging because they require the simultaneous analysis of genetic drift in several populations under various model assumptions. The demographic models analyzed with these methods are defined in terms of the parameters needed to describe the past growth of each population, their times of divergence from one another and the history of admixture among them.

Gutenkunst *et al* (2009) developed an efficient way to numerically solve a set of coupled diffusion equations and then search parameter space for the maximum likelihood parameter estimates. Their program *dadi* can analyze data from as many as three populations. Harris and Nielsen (2013) use the length distribution of tracts identical by descent within and between populations to estimate model parameters. Their program (unnamed) can handle the same degree of demographic complexity as *dadi*. Excoffier *et al* (2013) use coalescent simulations to generate the joint frequency spectra under specified demographic assumptions. Their program *fastsimcoal2* approximates the likelihood and then searches for the maximum likelihood estimates of the model parameters. Using simulations instead of numerical analysis allows *fastsimcoal2* to analyze a much larger range of demographic scenarios than *dadi*. Schiffels and Durbin (2014) recently introduced the multiple sequential Markovian coalescent (MSMC) model, which is a generalization of the pairwise sequential Markovian coalescent (PSMC) model (Li and Durbin 2011). MSMC uses the local heterozygosity of pairs of sequences to infer past effective population sizes and times of divergence.

These and similar methods are especially useful for human populations for which the historical and archaeological records strongly constrain the class of models to be considered. Although human history is much more complicated than tractable models can describe, those models can nonetheless reveal important features of human history that have shaped current patterns of genomic variation.

In this paper, we introduce another way to characterize genomic data from two or more populations. Our method is designed to indicate the past relationship between a single genome and one or more populations that have already been well studied. Our method is particularly useful for detecting small amounts of admixture between populations and the direction of that admixture, but it can also indicate the occurrence of bottlenecks in population size. Furthermore, it can also serve as a test of consistency with results obtained from other methods. We first introduce our method and apply it to models of two and three populations, focusing on the effects of gene flow and bottlenecks. Then we present the results of analyzing human and archaic hominin genomes. Some of the patterns in the data are consistent with simple model predictions and others are not. We explore specific examples in some detail in order to show how our method can be used in conjunction with others. Finally, we use projection analysis to test demographic inferences for European and Yoruba populations obtained from the four previous studies described above.

## Analytic theory

We assume that numerous individuals from a single population, which we call the *reference population*, have been sequenced. We also assume there is an outgroup that allows determination of the derived allele frequency, *x*, at every segregating site in the reference population. We next define the *projection* of another genome, which we call the *test genome*, onto the reference population. For each segregating site in the reference population, a weight *w* is assigned to that site in the test genome as follows. If the site is homozygous ancestral, *w*=0; if it is heterozygous, *w*=1/(2*x*); and if it is homozygous derived *w*=1/*x*. The projection is the average weight of sites in the test genome at which the frequency of the derived allele in the reference population is *x*, 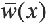.

With this definition of the projection, 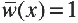 independently of *x* if the test genome is randomly sampled from the reference population. Therefore, deviation of 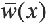 from 1 indicates that the test genome is from another population. To illustrate, assume that the test and reference populations have been of constant size *N*, that they diverged from each other at a time *τ* in the past, and that there has been no admixture between them since that time. The results of Chen *et al*. (2007) show that in this model 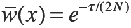 independently of *x*.

Analytic results are not so easily obtained for other models. We used numerical solutions to the coupled diffusion equations when possible and coalescent simulations when necessary to compute the projection under various assumptions about population history. For all models involving two or three populations, numerical solutions for each set of parameter values were obtained from *dadi* (Gutenkunst *et al* 2009). Models with more than three populations were simulated using *fastsimcoal2* (Excoffier *et al* 2013).

For all models we considered, an ancestral effective population size (*N*_*e*_) of 10000 with a generation time of 25 years was used. We assumed 150 individuals were sampled from the reference population and one from the test population. In *dadi* and *fastsimcoal2*, the resulting frequency spectrum was transformed into the projection for each frequency category. The parameters used are described in the figure captions.

### Two populations

We first consider two populations of constant size that separated *τ* generations in the past and experienced gene flow between them after their separation. We allow for two kinds of gene flow, a single pulse of admixture in which a fraction *f* of one population is replaced by immigrants from the other, and a prolonged period of gene flow during which a fraction *m* of the individuals in one population are replaced each generation by immigrants from the other. We allow for gene flow in each direction separately. Figure 1 shows typical results. Gene flow from the reference into the test population has no detectable effect while gene flow from the test into the reference results in the pattern shown: 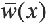 decreases monotonically to the value expected in the absence of gene flow. Even very slight gene flow in this direction creates the observed pattern. The projection is not able to distinguish between a single pulse and a prolonged period of gene flow, however. By adjusting the parameters, the projection under the two modes of gene flow can be made the same, as shown.

**Figure 1:**
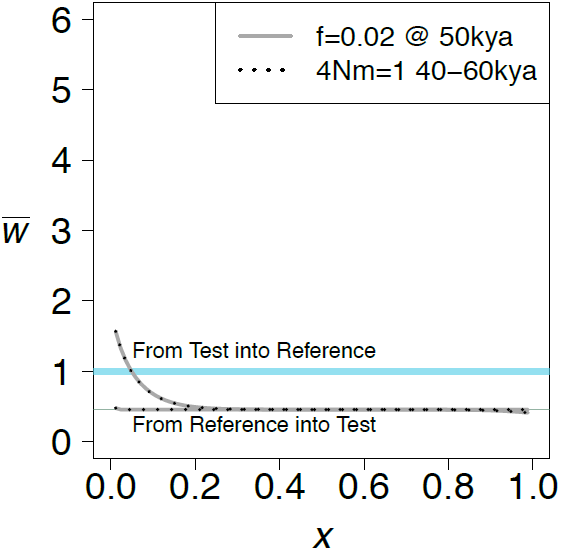
The effect of unidirectional gene flow on the projection of a test genome onto a reference population. Two kinds of gene flow were assumed, either a single pulse of admixture of strength *f*, or a period of immigration at a rate *m* per generation. Both populations are of constant size *N*=10,000. The divergence time, *τ*, is 400,000 years.

The intuitive explanation for the effect of gene flow from the test to the reference is that gene flow carries some alleles that were new mutations in the test population. Those alleles will necessarily be in low frequency in the reference population because they arrived by admixture but they are likely to be in higher frequency in the test population because they were carried by admixture to the reference. Therefore, they will be seen in the test genome more often than expected on the basis of their frequency in the reference population.

The projection changes from a horizontal line when there is a bottleneck in the reference (Fig. 2A, black line) or ancestral population (Fig. 2A, blue line) but not when there is a bottleneck in the test population (Fig. 2A, red line). The reason for the humped shape of the projection when there is a bottleneck in the reference population is that the bottleneck distorts the site frequency spectrum in that population in such a way that there are more rare and more common alleles than in a population of constant size and fewer alleles with intermediate frequency, and it accelerates the rate of loss of alleles that were previously in low frequency. When the reference population declines without recovering, the effect is an increase in rare alleles, similar to that of admixture into the reference population (Figure 2B, blue line). When the reference population expands, a slight decrease in rare alleles is observed (Figure 2B, red line).

**Figure 2:**
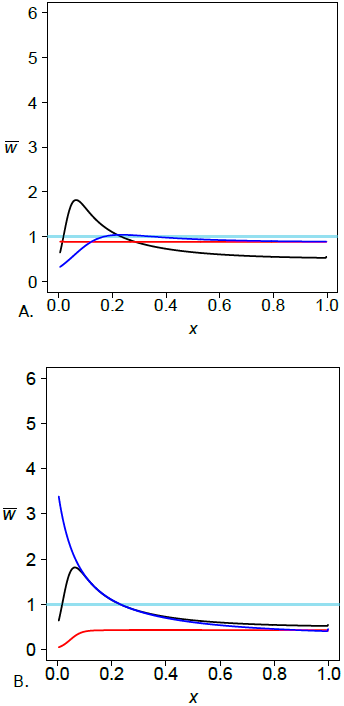
The effect of population size changes in a model with two populations diverged at time *τ*=60,000 years. In part A, a bottleneck occurs in the reference population (black), the test population (red), or the ancestral population (blue). During the bottleneck, the population size is reduced from 10,000 to 1,000 from 5020 kya. In part B, for the reference population only, a bottleneck occurs as in part A (black), a population expansion from 1,000 to 10,000 occurs 20 kya (red) or the reference population decreases in size from 10,000 to 1,000 50 kya (blue). The test population has the same population size as the ancestral population.

A bottleneck followed by admixture amplifies the effect of admixture (Figure 3A, black line) while admixture that occurs before or during the bottleneck does not change the shape of the projection as much (Figure 3A, red and blue lines). The effect comes from the increase in population size at the end of the bottleneck, not the decrease at the beginning (Fig. 3B).

**Figure 3:**
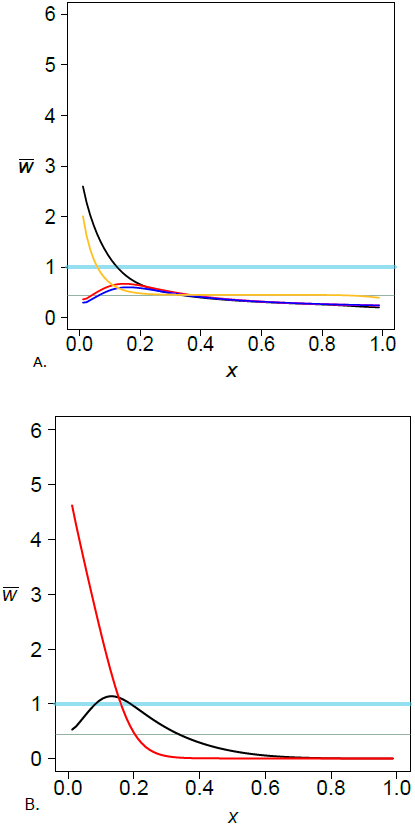
The combined effect of a bottleneck and admixture. The divergence time was *τ*=100,000 years. For A, the yellow projection represents no bottleneck but admixture of *f*=0.02 at 40 kya. The other projections include admixture at 40 kya (black), 80 kya (red), and 120 kya (blue) of 0.02 from the test to the reference, where there was a bottleneck from 70 – 90 kya. The bottleneck reduced the reference population size from 10,000 to 1,000, and it increased to 10,000. For B, the reference population size increased from 1,000 to 10,000 at 40 kya only. The divergence time was *τ*=100,000 years. Admixture of 0.02 from the test to the reference occurred at 30 kya (red) and 50 kya (black).

### Three populations

Three populations lead to a greater variety of effects than can be seen in two. Because samples are analyzed from only two of the populations, the test and the reference, the third population is unsampled. We will follow Beerli (2004) and call the unsampled population a *ghost population*. In some situations, all populations may be sampled but only two at a time are analyzed. In others situations, no samples are available from a population that is known or suspected to have admixed with one or more of the sampled populations. In the latter case, one goal is to determine whether or not there has been admixture from the ghost population.

We first consider the effects of gene flow alone. We will assume a single pulse of admixture of strength *f* at a time *t*_GF_. There are three distinct topologies representing the ancestry of the three populations (Fig. 4). Gene flow can be from the ghost population into either the test or the reference population. Gene flow from the ghost into the test population has little effect in all three topologies (Fig. 5A-C). How ghost gene flow into the reference population affects the projection depends on the population relationships. If the test and ghost populations are sister groups (Fig. 4A, 5D), then the effect is similar to that of gene flow directly from the test into the reference population (Fig. 1). When gene flow is directly from the test population, the higher value of 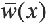 for small *x* reflects new mutations that arose in the test population after separation from the reference. When gene flow is from the ghost population, only mutations that arose in the internal branch contribute. Thus, when there is a longer period of shared ancestry between the test and ghost populations, there is a stronger effect of admixture (Fig. 5D).

**Figure 4:**
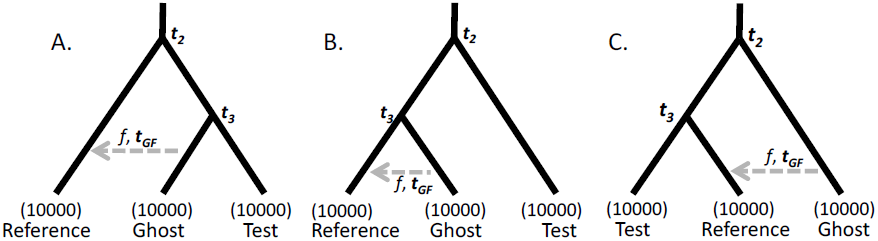
Illustration of three possible population relationships in which there is a pulse of admixture of intensity *f* at time *t*_GF_ in the past from the ghost population into the reference population. *t*_2_ and *t*_3_ are the times of population separation.

**Figure 5:**
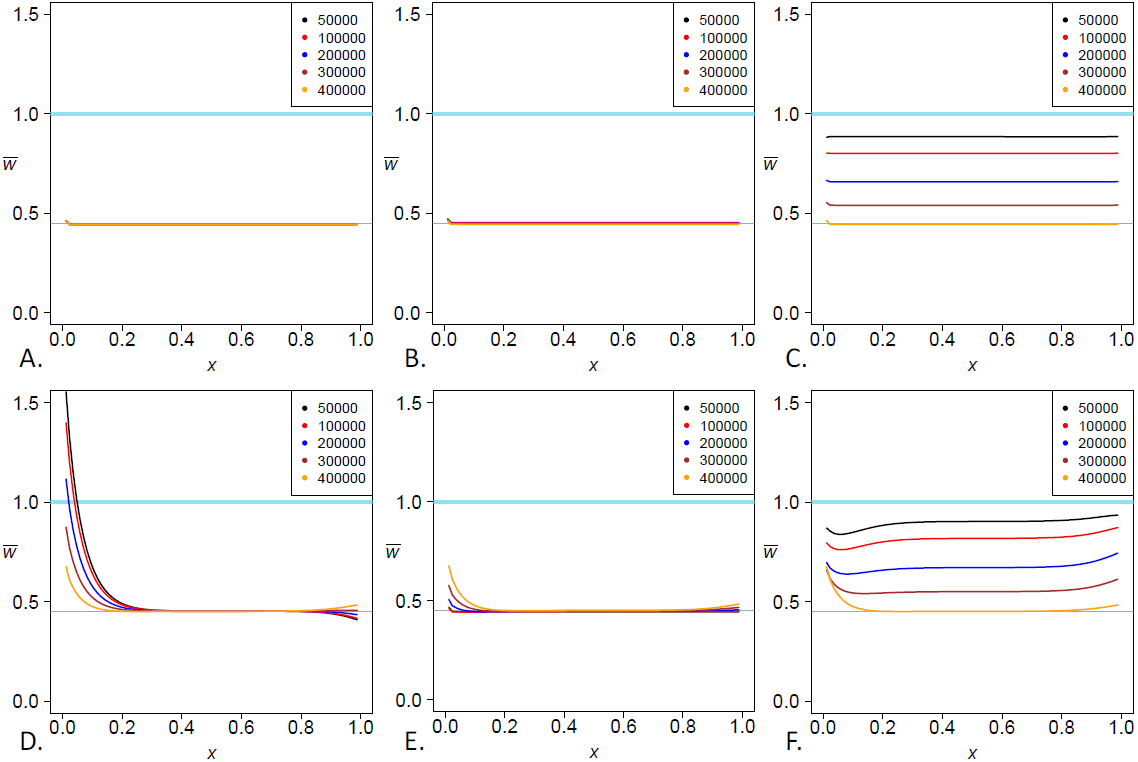
The effect of ghost admixture into the test (A-C) and the reference (D-F). A and D follow the topology in Fig. 4A, B and E follow Fig. 4B, and C and F follow Fig. 4C. *t*_2_ = 400 kya, *f* = 0.02, and *t*_*GF*_ = 50 kya. *t*_3_ is varied from 50 kya to 400 kya, according to the legend. Population sizes remain constant at 10,000.

In the second topology (Fig. 4B), the reference and ghost populations are sister groups. Here, gene flow from the ghost population has the opposite effect on the projection. For small *x*, the increase in 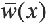 is reduced in magnitude when the shared ancestry between the reference and ghost populations increases. The intuitive reason is that some alleles carried from the ghost to the reference population arose as new mutations in the ghost population and hence cannot be in the test population. Therefore, there are fewer low frequency alleles in the test genome than expected. For a given time and magnitude of admixture, the effect increases as the time of separation of the ghost and reference populations increases (Fig. 5E). When the reference and test populations are sister groups (Fig. 4C), and the ghost population is an outgroup, a dip is observed for low frequencies and a slight increase is observed for common alleles (Fig. 5F).

If there is a bottleneck in the reference population after admixture, the effect (Fig. 6A) is similar to that seen in the two-population case (Fig. 3). The signal of admixture is amplified. In the case where the reference and ghost populations are sister groups (Fig. 6B), the characteristic bottleneck effect is observed. As the time of divergence between the reference and ghost population increases, the humped shape due to the bottleneck is reduced in size, presumably due to the increased effect of admixture. When the reference and test populations are sister groups, the humped shape remains, but the effect is reduced as the time of divergence increases (Fig. 6C), and the increase in common alleles is still observed.

**Figure 6:**
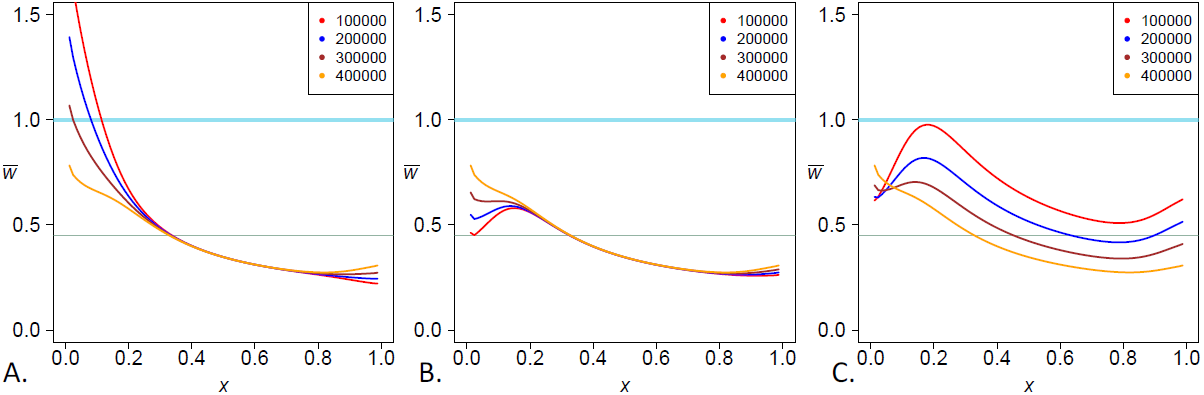
The effects of ghost admixture into the reference with a bottleneck in the reference occurring 70-100 kya changing the reference population size from 10,000 to 1,000 and back to 10,000. *t*_3_ is varied from 100 kya to 400 kya. All other parameters are the same as in Fig. 5.

### Ancestral Misidentification

Misidentification of the ancestral allele leads to the assumption that an allele is ancestral when it is in fact derived, or that an allele is derived when it is in fact ancestral. Hernandez *et al* (2007) show that ancestral misidentification occurs at levels of approximately 1-5% in human genome datasets. We use *ms* (Hudson 2002) to simulate two simple two-population projections to determine the effect of ancestral misidentification on the projection, one with no admixture or population size changes between the reference and test populations, and one matching the model with admixture shown in Fig. 1. We allowed for 0, 0.1%, 1% or 10% of the sites to be misidentified, reversing the ancestral or derived result given by the simulation. Where the frequency spectrum is shown to have an increase for common alleles (Hernandez *et al* 2007), the projection shows a similar result (Supplementary Fig. 1).

## Application to humans and archaic hominins

We will illustrate the use of projection analysis by applying it to genomic data from present-day humans and two archaic hominins (Neanderthal and Denisovan). For the reference populations, we used data from the 1000 Genomes (1000G) project for three populations, Europeans (CEU), Han Chinese (CHB) and Yoruba (YRI) (1000 Genomes Consortium 2010). For test genomes, we used the high-coverage Denisovan genome (Meyer *et al* 2012), the high-coverage Neanderthal genome (Prüfer *et al* 2014), and some of the high-coverage present-day human genomes sequenced by Meyer *et al* (2012). We will identify the reference populations by the 1000G abbreviation (CEU, CHB and YRI), and the test genomes by the names used by Meyer *et al* (2012). We used only autosomal biallelic sites with data present in every individual and population sampled. We used the reference chimpanzee genome, *PanTro2*, to determine the derived and ancestral allele at each site, and filtered out all CpG sites.

To show that projections give insight into human demographic history, we developed a ten-population demographic history with realistic parameters taken from the literature and adjusted using different curve-fitting techniques (Table 1, Fig. 7). The initial parameter ranges we chose were informed by a variety of previous studies, as noted in Table 1. To improve the fit of the simulated model to the projections, we used two techniques. Initially, we focused on two populations at a time. Using *dadi* (Gutenkunst *et al* 2009) and the Broyden-Fletcher-Goldfarb-Shanno algorithm (Morales and Nocedal 2011), we estimated several demographic parameters simultaneously that gave the best fitting projection for the two populations. For more than two populations, we used *fastsimcoal2* (Excoffier *et al* 2013) and Brent’s algorithm to vary one parameter at a time, fixing all other parameters. The parameters of interest were cycled through, each varied in turn, until a better fitting projection could not be found. This technique tended to converge most quickly when we focused on no more than three or four parameters at a time. For both techniques, we used the sum of least squares (LSS) to determine the best fit.

**Table 1:**
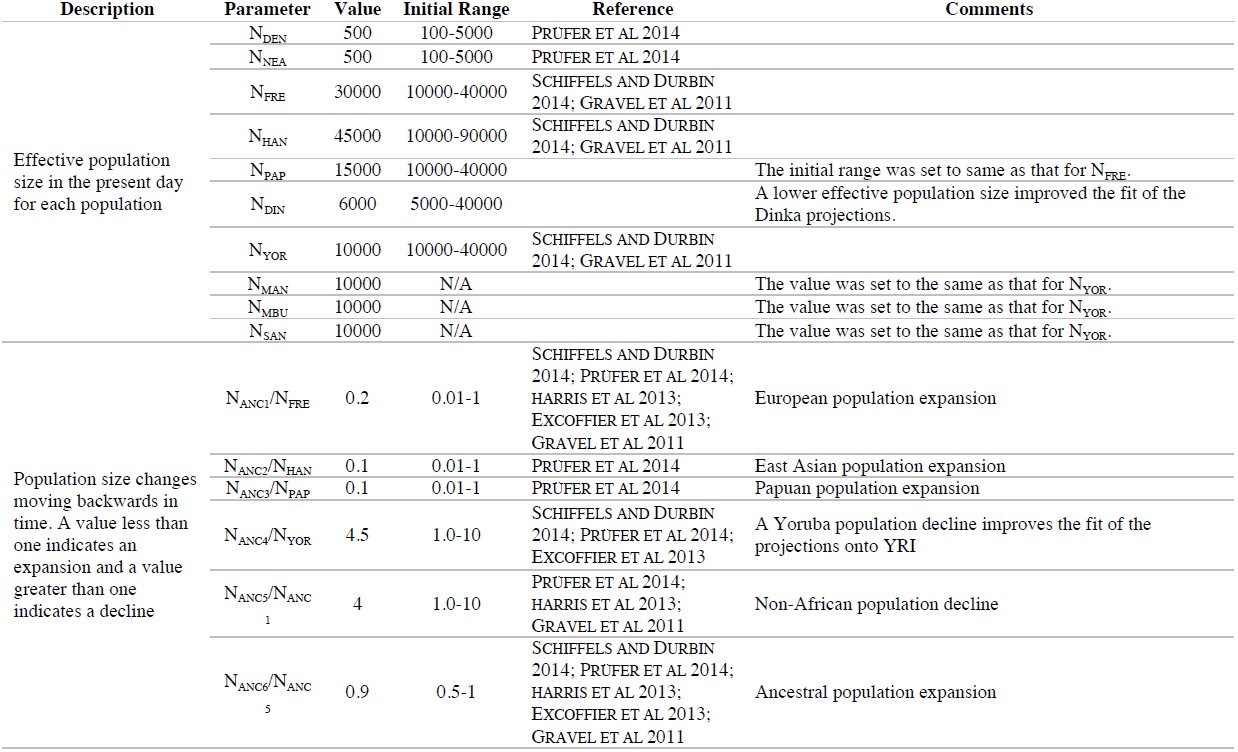
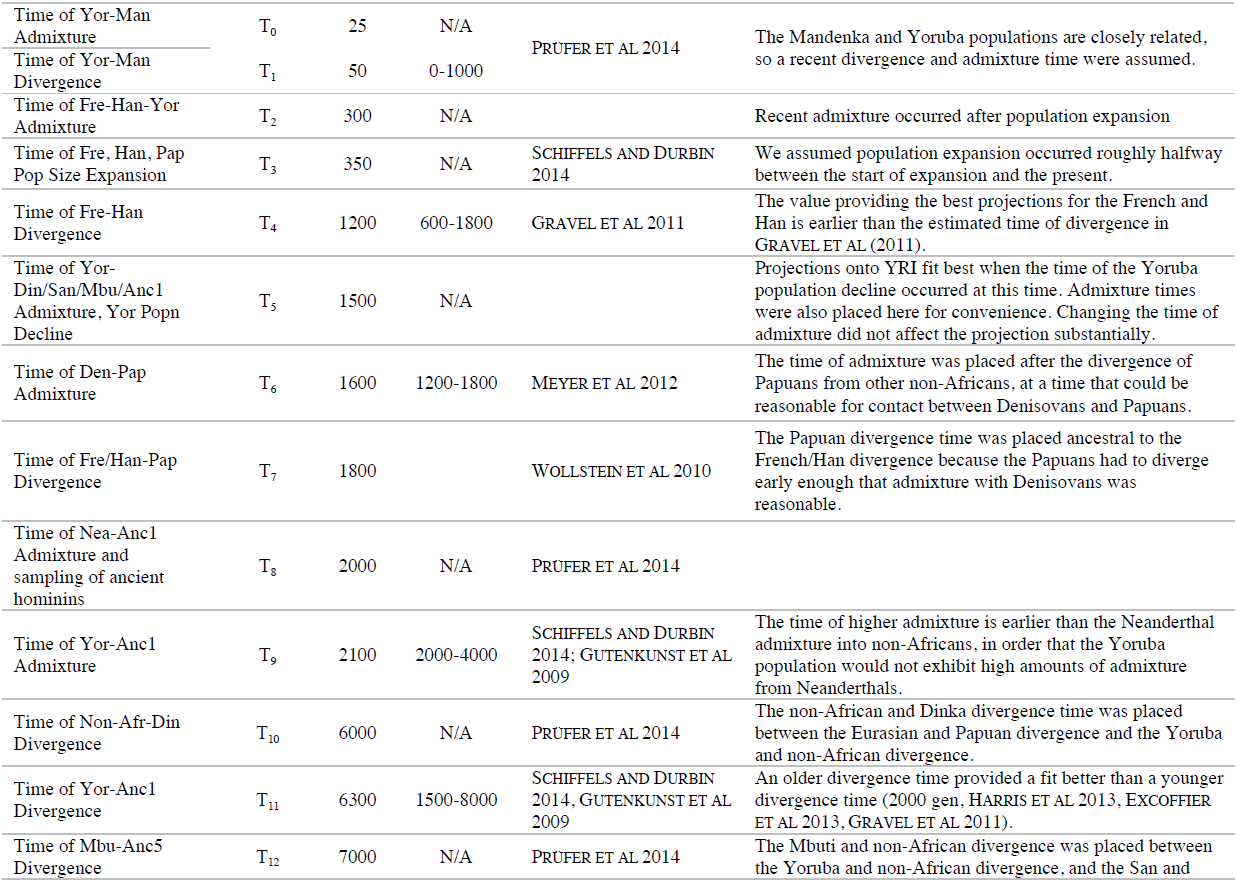
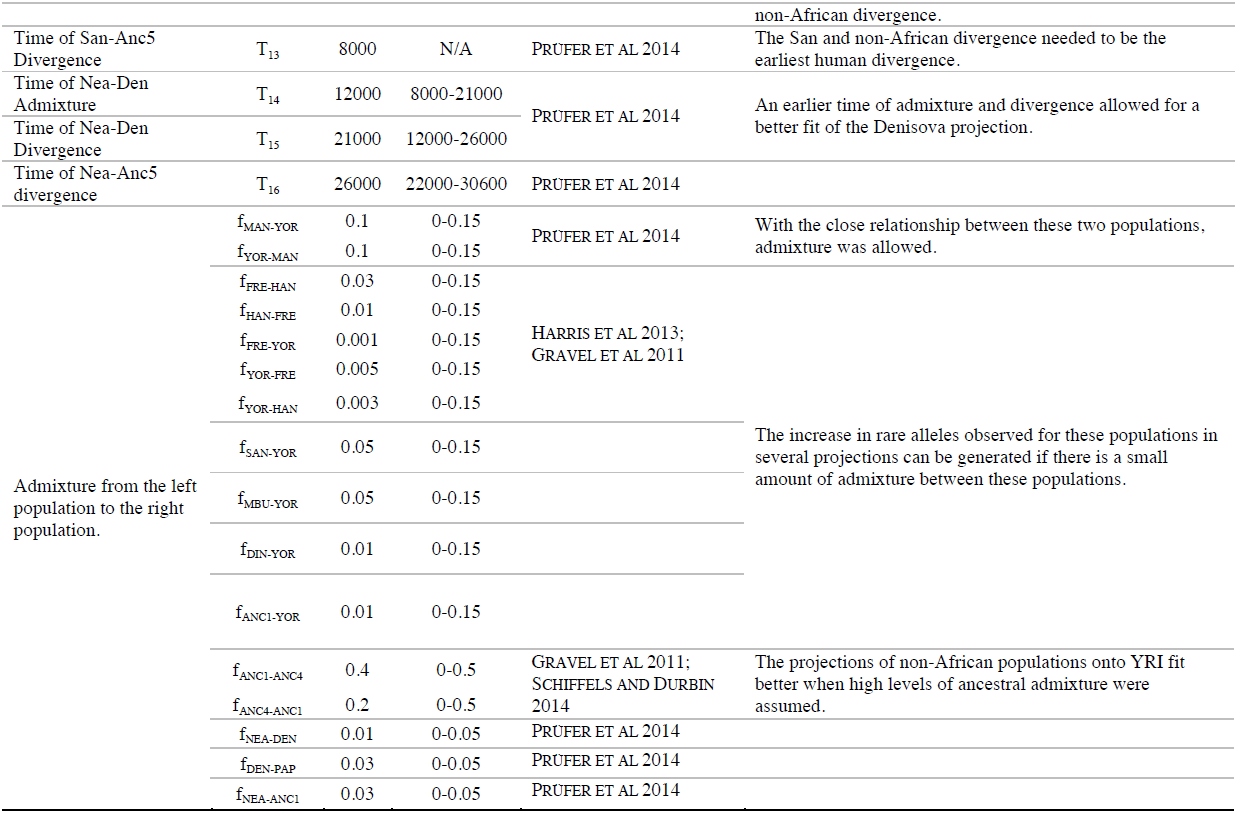
Description of parameters used in the simulation of the ten-population tree in Figure 7. The initial range is the set of values that was explored for each parameter. N/A indicates that the parameter was not varied. The initial range choices were based on the papers cited, though the ranges were sometimes expanded to explore the effects of more values. Times are in generations, with 1 gen = 25 years.

**Figure 7:**
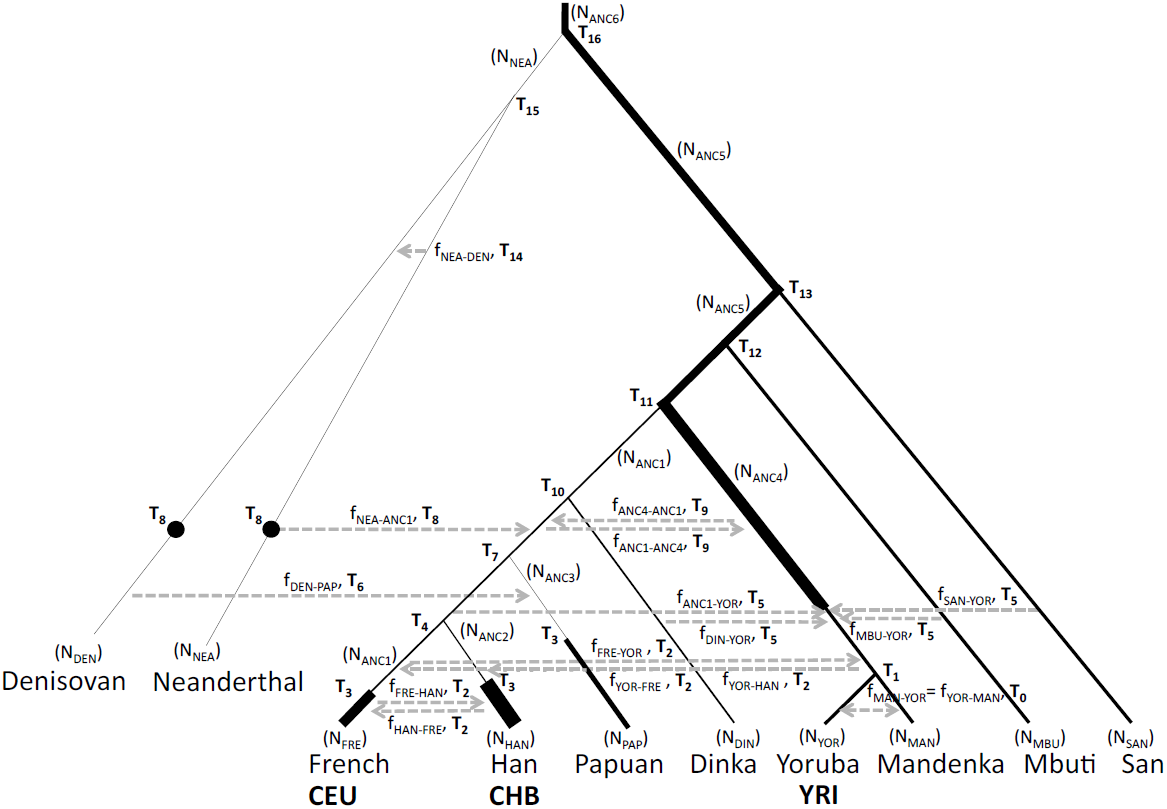
A model of human demographic history for ten populations that when simulated, gave projections similar to the observed projections. The bolded populations are the reference populations, and the row above them indicates the population origin of each test genome. The values used are found in Table 1. Black dots indicate the time of sampling if not in the present day. Thickness of the branch gives an approximation of the change in effective population size.

The demographic scenario displayed in Fig. 7 is not meant to be optimal. Instead, it is intended to show that, for a plausible scenario, the predicted projections are similar to ones computed from the data. This model illustrates the sensitivity of projections to major demographic processes that have shaped human history. Here, we note what features of demographic history are necessary to give rise to projections similar to those observed.

### Comparison of observed projections to each other

The black curves in Fig. 8-11 represent the observed projections. The projections were smoothed using a cubic spline and a smoothing parameter of 0.5. This was done to reduce the effect of sampling error in comparisons with the expected projections for the ten population demographic scenario described in Fig. 7, which are represented by the red curves. Supplementary Tables 1-3 provide the LSS comparing the projections of each test genome onto each reference population, and the diagonal terms provide the LSS for that test genome, relative to the 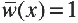 line. The observed projections show that the Neanderthal and Denisovan projections onto CEU, CHB and YRI look the most different from the 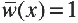 line.

**Figure 8:**
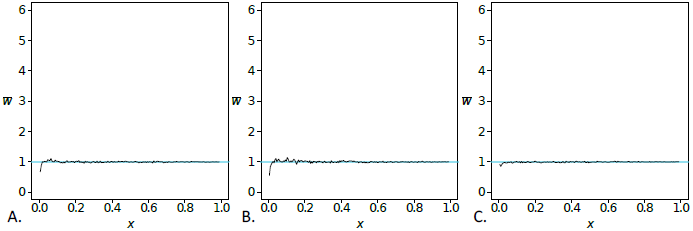
The projections of French onto CEU (A), Han onto CHB (B), and Yoruba onto YRI (C). The sum of least squares (LSS) scores comparing the observed projection to the 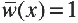 line are found in Supplementary Tables 1-3.

### Comparison of a test genome with the same population

In Fig. 8A, the projection of the French genome onto CEU fits the expectation except for small *x*. Similar deviations are seen in Fig. 8B in the projection of the Han genome onto CHB and, to a lesser extent, in Fig. 8C in the projection of the Yoruba genome onto YRI. This pattern is expected for the smallest frequency classes because the frequency spectrum in the reference populations has more singletons than expected in a population at equilibrium under drift and mutation. See the Appendix for details.

### Admixture with Neanderthals and Denisovans

Our simulations show that a bottleneck combined with admixture into the reference population can result in a strong effect on the projection (Fig. 3A, black curve). The projections of the Altai Neanderthal onto CEU and CHB show a large excess of rare alleles (Fig. 9I and 10I), which requires the combination of a bottleneck in the ancestors of non-Africans and admixture from Neanderthals into non-Africans after that bottleneck. Including both processes in our model, we obtain good fits to the observed projections (Table 2, Fig. 9I and 10I). When either admixture or the bottleneck is omitted, the result is a decrease in the excess of rare alleles and a worse fit (Supplementary Table 4, Supplementary Fig. 2).

**Figure 9:**
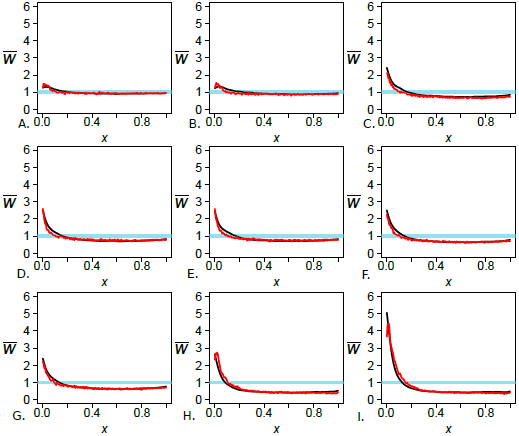
The observed projection (black) and simulated projection from our model (red) for the CEU reference population. The test genomes are Han (A), Papuan (B), Dinka (C), Yoruba (D), Mandenka (E), Mbuti (F), San (G), Denisovan (H), and Neanderthal (I). The LSS scores comparing the observed projections to each other and 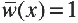 line can be found in Supplementary Table 1, and the LSS scores comparing the observed and simulated projections can be found in Table 2.

**Figure 10:**
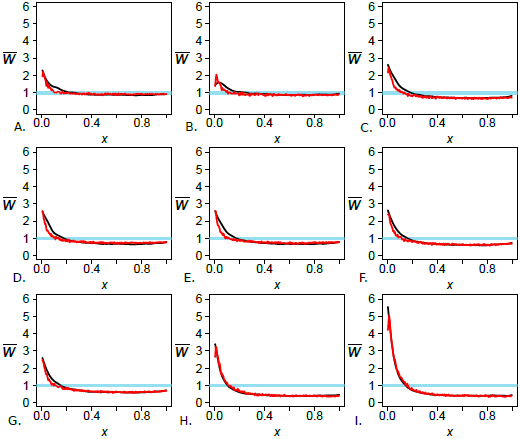
The observed projection (black) and simulated projection from our model (red) for the CHB reference population. The test genomes are French (A), Papuan (B), Dinka (C), Yoruba (D), Mandenka (E), Mbuti (F), San (G), Denisovan (H), and Neanderthal (I). The LSS scores comparing the observed projections to each other and 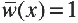 line can be found in Supplementary Table 2, and the LSS scores comparing the observed and simulated projections can be found in Table 2.

Similarly, the projections of the Denisovan genome onto CEU and CHB (Fig. 9H and 10H) are consistent with the three-population analysis shown in Fig. 5D. In this case, Neanderthals are the ghost population and Denisovans are the test population. The excess of rare alleles for the Denisova projection is consistent with Neanderthals and Denisovans being sister groups. Some of the new mutations that arose in the shared branch between Neanderthals and Denisovans are carried by admixture to humans and their presence is seen in the projection. When the Denisova and Neanderthal populations are sister groups, there is an excess of rare alleles that is predicted by our model (Table 2, Fig. 9H and 10H). The Denisovan projections give a signal of admixture but it is weaker than the signal in the Neanderthal projections.

The projections of the Neanderthal (Fig. 11I) and Denisovan (Fig. 11H) onto YRI show a signal of admixture even though previous analysis of the Neanderthal genome did not find evidence of direct Neanderthal admixture from the presence of identifiable admixed fragments (Prüfer *et al* 2014). These projections are consistent with the signal of Neanderthal introgression being carried by recent admixture from the ancestors of CEU and CHB into the ancestors of YRI. In our model (Fig. 7), there is no admixture between YRI and any archaic hominin, but there is gene flow between the ancestors of YRI and non-Africans. An excess of rare alleles is observed in the simulated projection (Fig. 11H and 11I). Admixture from non-Africans to Yoruba had to have occurred more recently than the Neanderthal admixture into non-African populations for this signal to be present.

**Figure 11:**
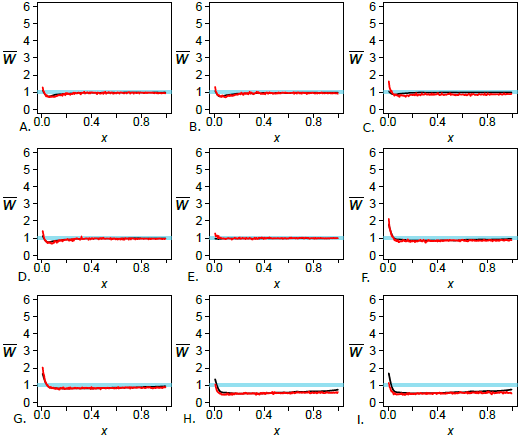
The observed projection (black) and simulated projection from our model (red) for the YRI reference population. The test genomes are Han (A), Papuan (B), Dinka (C), French (D), Mandenka (E), Mbuti (F), San (G), Denisovan (H), and Neanderthal (I). The LSS scores comparing the observed projections to each other and 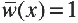 line can be found in Supplementary Table 3, and the LSS scores comparing the observed and simulated projections can be found in Table 2.

The Altai Neanderthal genome is unusual in that it is marked by long runs of homozygosity, indicating the individual was highly inbred. Prüfer *et al* (2014) show that the inbreeding coefficient was 1/8. This inbreeding has no effect on the projection, however, because the projection effectively samples a haploid genome from the test individual.

### Relationship among Non-African Populations

The projection of the French genome onto CHB (Fig. 10A) differs from the projection of the Han onto CEU (Fig. 9A). This difference reflects the subtle interplay between admixture and population size changes. A model in which the ancestors of CHB experienced a bottleneck after their separation from the ancestors of CEU along with a greater rate of population expansion can explain why the humped shape characteristic of bottlenecks was not swamped out by the signal of admixture. The inclusion of more admixture from CEU to CHB than the reverse can account for the overall increased excess seen in the French projection onto CHB (Fig. 10A). When these events are included in our model, the resulting projections are relatively close to the observed projections (Table 2).

The Papuan demographic history modeled here includes divergence from the ancestors of Europeans and East Asians and a bottleneck and population expansion (Fig. 7). In this model, we simulated a demographic history in which the Papuans diverged from the population ancestral to Europeans and East Asians, a scenario supported by Wollstein et al (2010), but not by others (Prüfer *et al* 2014, Meyer *et al* 2012). We made this assumption because we followed (Gravel *et al* 2011) in assuming that Europeans and East Asians diverged relatively recently. With admixture from Denisovans to Papuans occuring earlier, assuming the Papuans were the outgroup to Europeans and East Asians was more appropriate. Using this model, the projections fit relatively well (Table 2, Fig. 9B and 10B).

### Relationship Between Non-Africans and YRI

The projections of the Papuan, French and Han genomes onto YRI (Fig. 11A, 11B, and 11D) are similar despite the difference between the Han and Papuan projections onto CEU (Fig. 9A and 9B). These observations can be accounted for if there were high levels of admixture between the ancestors of non-Africans and the ancestors of the Yoruba population as well as a large ancestral Yoruba population that declined in the recent past. These two processes together explain the dip observed and the increase to 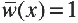 for larger *x*, and they lead to a good fit to the observed projections (Table 2, Fig. 11A, 11B, 11D). Varying these two parameters in our model shows their effect on the projection for rare alleles and that higher values for both of these parameters give the best fitting simulated projections (Supplementary Table 5, Supplementary Fig. 3).

### African projections onto CEU and YRI

The projections of all five African genomes—San, Yoruba, Mandenka, Dinka, and Mbuti—onto CEU (Fig. 9C-G) are similar to one another and similar to their projections onto CHB (Fig. 10C-G). All these projections are consistent with low levels of admixture from the African populations into the ancestors of CEU and CHB. This conclusion is surprising. Previous analyses (Meyer *et al* 2012, Prüfer *et al* 2014, Lachance *et al* 2012, Pickrell *et al* 2012) showed that the San population diverged from other African populations before the other African populations diverged from one another and before the ancestors of CEU and CHB diverged from each other. The separate history of the San is not reflected in their projection of the San genome onto CEU and CHB. Because the demographic history in the reference populations has a strong effect on the projections, the bottleneck in Europeans combined with low amounts of admixture between the Yoruba and San, and between the Yoruba and non-Africans are enough to give results similar to the observed projections (Table 2, Fig. 9C-G). A closer look at the middle of the projection for reference CEU shows that the San projection is slightly lower than the Yoruba projection (Fig. 9D and 9G), which suggests that the difference in divergence time is weakly reflected in the projection.

The projections of different African genomes (Dinka, Mandenka, Mbuti, San) onto YRI (Fig. 11C, 11E-G) illuminate the relationship between these four African populations and the Yoruba. Other studies (Prüfer *et al* 2014, Meyer *et al* 2012, Tishkoff *et al* 2009) have shown that, while the San and Mbuti are the most diverged from all other populations sampled, the Mandenka and Yoruba populations have only recently separated and the Dinka population shares some ancestry with non-African populations. The San and Mbuti projections onto YRI show a slight excess of rare alleles, suggesting some admixture from their ancestors into the ancestors of YRI. The Mbuti is closer to the 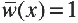 line, which suggests it is less diverged from YRI than is the San, agreeing with the model proposed in other studies (Prüfer *et al* 2014, Meyer *et al* 2012, Tishkoff *et al* 2009). The Mandenka projection falls nearly on the 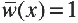 line, suggesting it is indistinguishable from a random YRI individual. Finally, the Dinka projection onto YRI exhibits a dip that is similar, though of reduced magnitude, to those observed in all the non-African projections, perhaps due to greater admixture between the ancestors of the Dinka and Yoruba in Africa. Including these events in the model (Fig. 7) gives a close fit to the observed projections (Table 2, Fig. 11C, 11E-G).

## Test of Published Models

We used observed projections to test for consistency with inferred demographic parameters from four studies (Gravel *et al* 2011; Excoffier *et al* 2013; Harris and Nielsen 2013; Schiffels and Durbin 2014) for European and Yoruba populations. All four studies applied their methods to CEU and YRI.

We obtained projections by using *fastsimcoal2* (Excoffier *et al* 2013) to simulate one million SNPs with the estimated demographic parameters from each of these four models. The demographic parameters used are shown in Figure 12. We compare the simulated projections to the observed projections of a Yoruba genome projected onto CEU and of a French individual onto YRI. The visual differences highlight aspects of each model that agree or disagree with the observed projections.

**Figure 12:**
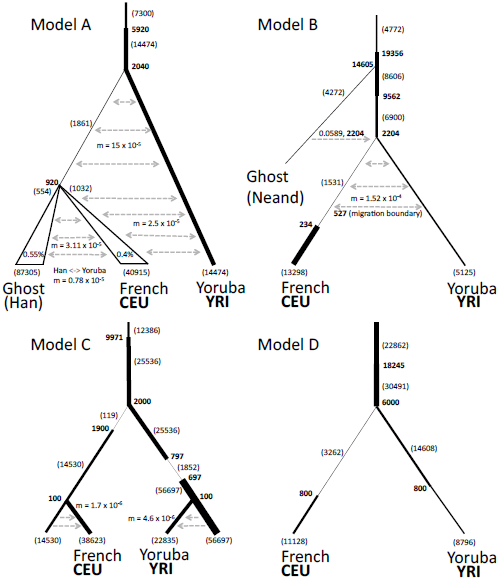
The demographic models from each of the four previous studies, Gravel *et al*. (2011, Model A), Harris and Nielsen (2013, Model B), Excoffier et al. (2013, Model C), and Schiffels and Durbin (2014, Model D). Shading and symbols have the same meaning as in Figure 7, and the triangle indicates growth at the given percentage.

The four models overlap but differ in the estimates of a number of parameters. All models assume a population decrease in ancestral Europeans, presumably during dispersal out of Africa. The severity of the population size change ranges from 0.0047 (Model C) to 0.22 (Model B) and occurs at the time when the ancestors of YRI and CEU diverged. Models A, B, and D assume a subsequent population expansion, while Model C, which has the most extreme reduction, recovers 100 generations after the population decrease. In Model A, YRI is assumed to be of constant size while the size declines in Models B and D. In Model C, the ancestral YRI population undergoes a bottleneck 797 generations ago. In all four models, the population ancestral to CEU and YRI increases in size before the two populations separated. In Models A, B and C, the time of divergence of CEU and YRI is about 50 kya. In Model D the separation time is at least 150 kya.

Model A assumes higher rates of migration soon after the CEU-YRI divergence and a lower rate more recently. Model B allows for migration between CEU and YRI and it also includes a parameter for ghost admixture from an archaic hominin that diverged 14605 generations ago (365 kya). Model C uses a continent-island model, in which CEU and YRI diverge from continental European and African populations recently, receiving migrants from those populations until the present. However, neither they nor their ancestral populations admix with each other. Model D does not allow for migration between CEU and YRI, though Schiffels and Durbin (2014) say that such migration probably occurred.

The simulated projections show that Model A gives the best fit to the observed projections (Table 3, Figure 13). For Model A, increasing the rate of recent migration from YRI to CEU from 0.000025 migrants/generation to 0.00005 migrants/generation led to a slightly better fit (Table 3, Figure 14). The other three models do not give projections that fit as well. Modifying the amount of admixture and/or migration in each of Models B-D resulted in a substantially better fit (Table 3, Figure 14). In Model B, increasing the migration rate from YRI to CEU and adding admixture 150 generations ago at a rate of 0.02 from CEU to YRI and a reverse rate of 0.015 resulted in a better fit. In Model C, adding admixture at two different times led to a better fit. We first added recent admixture at a rate of 0.07 150 generations ago from Europeans to Yoruba with a reverse rate of 0.1. Then, we added ancestral admixture at a rate of 0.37 from Europeans to Yoruba and a reverse rate of 0.2 1710 generations ago. In Model D, adding symmetric admixture of 0.01 150 generations ago between Yoruba and Europeans, and allowing for migration beginning at 1662 generations ago of 0.0007 migrants/generation from Europeans to Yoruba and 0.0003 migrants/generation from Yoruba to Europeans results in a better fit (Table 3, Models A*-D*, Fig. 14).

**Figure 13:**
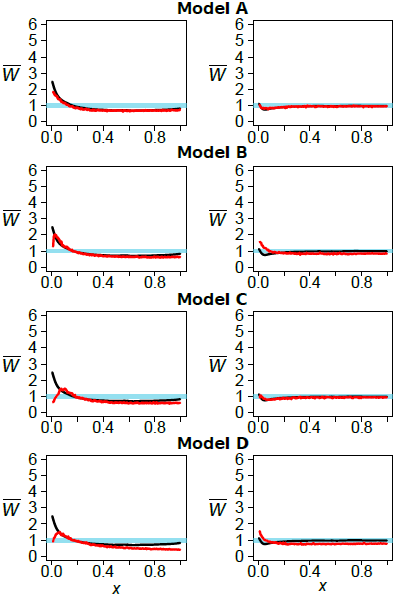
The observed projections (black) and simulated projections from demographic models inferred from other studies (red). The left column is the Yoruba genome projected on CEU and the right column is the French genome projected on YRI. The rows represent the estimates from Models AYD. LSS scores are in Table 3.

**Figure 14:**
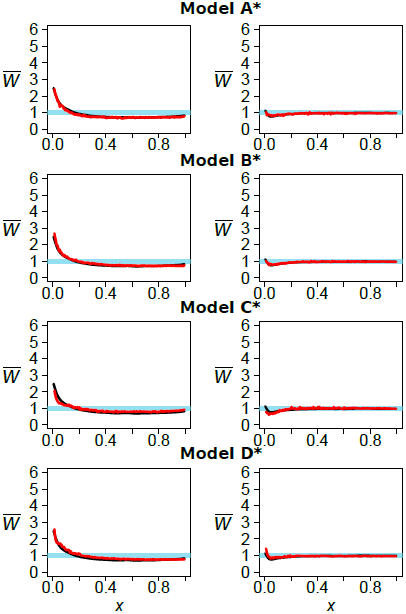
Projections for previous studies (Models A-D) where the parameters for migration or admixture between Europeans and Yorubans have been added or modified for a better fit. The left column is the Yoruba genome projected onto CEU and the right column is the French genome projected onto YRI. LSS scores are in Table 3.

Our projection analysis supports the hypothesis that there was significant gene flow between the ancestors of CEU and YRI after there was introgression from Neanderthals into CEU. Gravel *et al* (2011) (Model A) reached this conclusion by allowing for such gene flow in their analysis. Adding such gene flow to Models B-D substantially improved the fits to the observed projections.

## Discussion and Conclusions

We have introduced projection analysis as a visual way of comparing a single genomic sequence with one or more reference populations. The projection summarizes information from the joint site frequency spectrum of two populations. We have shown that projections are affected by various demographic events, particularly population size changes in the reference population and admixture into the reference population. The time since two populations had a common ancestor also affects the projection, as does the interaction with unsampled populations.

Projection analysis is primarily a visual tool and is not intended to replace methods such as those developed by Gutenkunst *et al* (2009), Harris and Nielsen (2013), Excoffier *et al* (2013), and Schiffels and Durbin (2014) that estimate model parameters. Projection analysis uses less information than these methods. Instead projection analysis is intended to be a method of exploratory data analysis. It provides a way to compare a single genomic sequence, perhaps of unknown provenance, with several reference populations, and it provides a way to test the consistency of hypotheses generated by other means.

Our applications of projection analysis to human and archaic hominin populations largely confirmed conclusions from previous studies. In particular, we support the hypothesis that Neanderthals admixed with the ancestors of Europeans and Han Chinese and the hypothesis that Neanderthals and Denisovans are sister groups.

By analyzing present day human populations, we provide strong support for the conclusion of Gutenkunst *et al* (2009) and Gravel *et al* (2011) that there was continuing gene flow between the ancestors of Yoruba and the ancestors of Europeans long after their initial separation. The fit of other models improves when such gene flow is included.

Harris and Nielsen (2013) incorporate migration in their model, but they assume a small amount from the time of separation until a few thousand years ago. The Excoffier *et al* (2013) model does provide a good fit for the French projection onto YRI, perhaps because of the large bottleneck they infer in the ancestral Yoruba, but the Yoruba projection onto CEU requires some admixture for a better fit. Schiffels and Durbin’s (2014) model does not allow for estimation of migration parameters. However, they argue that there was probably an initial divergence with subsequent migration before a full separation. Our conclusion is consistent with theirs. There was likely substantial gene flow between the ancestors of Europeans and Yoruba after their initial separation but before movement out of Africa. Then, stronger geographic barriers led to lower rates of gene flow and effectively complete isolation.

Throughout we have assumed that population history can be represented by a phylogenetic tree. Although that assumption is convenient and is made in most other studies as well, we recognize that a population tree may not be a good representation of the actual history. For example, the inferred period of gene flow between Europeans and Yoruba may actually reflect a complex pattern of isolation by distance combined with the appearance and disappearance of geographic barriers to gene flow. At this point, introducing a more complex model with more parameters will not help because there is insufficient power to estimate those parameters or to distinguish among several plausible historical scenarios.

The effect of ancestral misidentification on projection analysis was also a concern. Here, we show that low levels of ancestral misidentification lead to an increase in common alleles. Thus, we expect and do see a slight increase of 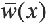 in common alleles in most observed projections.

Projection analysis is designed for analyzing whole genome sequences but it can be applied to other data sets including partial genomic sequences, dense sets of SNPs and whole exome sequences. However, ascertainment of SNPs could create a problem by reducing the sample sizes of low and high frequency alleles. Of course the smaller the number of segregating sites in the reference genome, the larger will be the sampling error in the projection. The number of samples from the reference population also affects the utility of the projection. As we have shown, an important feature of many projections is the dependence of 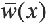 on small *x*. Relatively large samples from the reference population (50 or more individuals) are needed to see that dependence clearly. When sufficiently large samples are available, projection analysis provides a convenient way to summarize the joint site frequency spectra of multiple populations and to compare observations with expectations from various models of population history.

## Acknowledgements

This research was supported in part by a National Institutes of Health NRSA Traineeship (T32 HG 00047) and a National Science Foundation Graduate Research Fellowship Program Division of Graduate Education (1106400) to M.A.Y, and in part by a National Institutes of Health grant (R01-GM40282) to M.S. We thank N. Patterson, B. Peter, F. Racimo, D. R. Reich and J. G. Schraiber for helpful discussions of this topic and for comments on previous versions of this paper.

## Appendix: The projection of a test genome onto a reference population and applications to humans and archaic hominids

The aim of this appendix is to present a theoretical justification for the “dip” at low frequencies observed in Figure 8, which shows a French genome projected onto a CEU panel, a Chinese genome projected onto a Chinese panel, and a Yoruban genome projected onto a Yoruban panel. In each case, the test genome appears to carry fewer of the panel’s derived singletons and doubletons than expected given the close relationship between the test and reference genomes. We argue that this is a consequence of finite reference panel size in a species that has inflated counts of low frequency alleles due to recent population growth.

Each comparison in Figure 8 is akin to the scenario of starting with a population reference panel of *N* + 1 genomes, picking one genome uniformly at random, and projecting this “test” genome onto the remaining *N*-genome panel. If we fix a frequency *f* and let *N* go to infinity, it is trivial to see that the projection 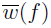 should approach *f*. However, this does not imply that 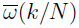 should equal *k*/*N* for *k* and *N* fixed.

We can compute the expected value of 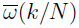 in terms of the frequency spectrum (*f*_1_, *f*_2_, …, *f*_*N*+1_) of the entire population sample, where *f*_1_ is the frequency of singletons, *f*_2_ is the frequency of doubletons, and so on. In terms of these frequencies, 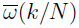 has the following expected value:

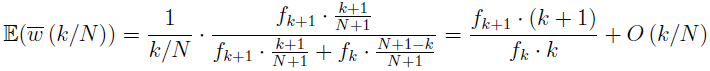

Here, the factor (*k* + 1)/(*N* +1) is the probability that the test individual has the derived allele given that *k* + 1 out of the *N* + 1 members of the original panel have the derived allele. Likewise, (*N* + 1 − *k*)/(*N* + 1) is the probability that the test individual has the ancestral allele given that *k* out of *N* + 1 panel members have the derived allele. This implies 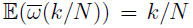 if and only if

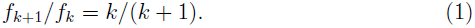

In a panmictic population that has reached effective population size equilibrium, coalescent theory does predict that *f*_*k*+1_/*f*_*k*_ = *k*/(*k* + 1). However, the site frequency spectrum is so sensitive to past changes in effective population size that equation (1) does not often hold for real datasets, and in general, low frequency variants show the most deviation from (1). In addition, some 1000 Genomes reference panel “singletons” may be sequencing errors that have a very low probability of being observed in a test genome because they are not true segregating genetic variants. Somatic cell line mutations are similarly unlikely to be shared. Cryptic population structure may be another source of deviation from the 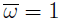 expectation at low allele frequencies.

Table 1 lists values of *W*_*k*_ := *f*_*k*+1_ · (*k* + 1)/(*f*_*k*_ · *k*) for the CEU, CHB, and YRI panels of the 1000 Genomes data, letting *k* range from 1 to 9. Assuming that the panel contains no sequencing errors, *W*_*k*_ is the expected value of 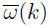 for projection of a same-population sequence onto the panel. Both the European and Chinese reference panels have *W*_*k*_ values that are less than 1 for *k* < 5 as a result of recent population growth, explaining the pronounced dip we see in these projections. In contrast, the YRI panel does not contain excess low frequency variants, suggesting that the smaller dip at *k* = 1 seen in the Yoruba projection may result from other causes such as sequencing error or reference panel population structure.

**Table 1:** Expected projection values *W*_*k*_ = *f*_*k*+1_ · (*k* + 1)/(*f*_*k*_ · *k*) for small values of *k* in three 1000 Genomes reference panels.

| Panel | $k = 1$ | 2 | 3 | 4 | 5 | 6 | 7 | 8 | 9 |
| --- | --- | --- | --- | --- | --- | --- | --- | --- | --- |
| CEU | 0.743 | 0.902 | 0.964 | 0.982 | 0.986 | 1.004 | 1.014 | 1.015 | 1.006 |
| CHB | 0.64 | 0.805 | 0.942 | 0.968 | 1.009 | 0.988 | 1.005 | 0.999 | 1.028 |
| YRI | 1.219 | 0.996 | 0.995 | 0.991 | 0.991 | 0.983 | 0.985 | 0.982 | 1.001 |

